# LuxS Inactivation Increases Antibiotic Susceptibility and Impairs Genetic Competence in *Streptococcus pneumoniae*

**DOI:** 10.64898/2026.11.17.767261

**Authors:** Gabriela Salvadori, Ilangko Balasingham, Fernanda C. Petersen

**Author notes:** Correspondence: Gabriela Salvadori.

## Abstract

The LuxS enzyme participates in bacterial metabolism and produces the precursor of autoinducer-2 (AI-2), a signaling molecule implicated in bacterial stress responses and genetic competence. Although LuxS/AI-2 has been associated with antibiotic responses in other streptococci, its contribution to antibiotic susceptibility in *Streptococcus pneumoniae* remains poorly understood. Here, we investigated the role of LuxS in pneumococcal survival during exposure to erythromycin and ampicillin and examined whether supplementation with 4,5-dihydroxy-2,3-pentanedione (DPD), the precursor of AI-2, could alleviate the effects of *luxS* inactivation. The *ΔluxS* mutant exhibited significantly greater reductions in viable cell counts than the wild type following antibiotic exposure, even after accounting for differences in baseline growth. Supplementation with 50 nM DPD significantly improved mutant survival during exposure to both antibiotics. We further investigated the contribution of LuxS/AI-2 to genetic competence, a phenotype previously associated with *luxS* inactivation. The mutant exhibited reduced transformation efficiency, which was significantly improved by DPD supplementation, although recovery remained partial. Together, these findings provide evidence that LuxS contributes to pneumococcal survival during antibiotic exposure and demonstrate that exogenous DPD can partially compensate for the effects of *luxS* inactivation on antibiotic responses and genetic transformation. These results extend our understanding of LuxS/AI-2-associated phenotypes in *S. pneumoniae*.

## INTRODUCTION

LuxS participates in cellular metabolism and produces the precursor of the quorum-sensing signal autoinducer-2 (AI-2). As part of the activated methyl cycle, this enzyme catalyzes the conversion of S-ribosylhomocysteine into homocysteine and 4,5-dihydroxy-2,3-pentanedione (DPD). DPD spontaneously undergoes chemical rearrangements to form the molecules collectively referred to as AI-2, which mediate communication in several Gram-positive and Gram-negative bacteria (Schauder et al., 2001; Miller et al., 2004). Because the same enzymatic reaction contributes to metabolism and signal production, phenotypes associated with *luxS* inactivation cannot readily be attributed to either function alone (Winzer et al., 2002).

The LuxS/AI-2 system has been implicated in streptococcal responses to antibiotic exposure. In *Streptococcus anginosus, luxS* inactivation increased susceptibility to ampicillin and erythromycin, and DPD supplementation restored growth during antibiotic exposure to levels similar to those of the wild type (Ahmed et al., 2007). In *Streptococcus intermedius*, LuxS/AI-2 affected biofilm formation and cell viability during exposure to subinhibitory antibiotic concentrations, and DPD supplementation complemented the mutant phenotype (Ahmed et al., 2009). These findings suggest that extracellular DPD/AI-2 can contribute to antibiotic stress responses in streptococci.

*Streptococcus pneumoniae* commonly colonizes the human nasopharynx asymptomatically but can cause diseases including otitis media, pneumonia, meningitis and bacteremia. Previous studies have linked pneumococcal LuxS to nasopharyngeal persistence, biofilm formation, carbohydrate utilization and virulence (Joyce et al., 2004; Trappetti et al., 2011, 2017). A connection with antibiotic stress was reported by Rogers et al. (2007), who observed increased *luxS* expression following exposure to a subinhibitory concentration of penicillin. This association raises the question of whether LuxS contributes to pneumococcal survival during antibiotic exposure. However, this functional contribution has not, to our knowledge, been directly investigated.

Beyond antibiotic stress, LuxS has been linked to pneumococcal genetic competence, the ability to take up extracellular DNA and incorporate it into the genome. Competence is controlled by a signaling pathway involving the competence-stimulating peptide (CSP). Encoded by *comC*, CSP is processed and exported by ComAB and activates the membrane receptor ComD. ComD then phosphorylates the response regulator ComE, which promotes expression of early competence genes, including those encoding the alternative sigma factor SigX, also known as ComX (Pestova et al., 1996; Lee and Morrison, 1999). SigX activates genes required for DNA uptake, processing and recombination (Lee and Morrison, 1999). Competence development integrates cell density, environmental conditions and cell history, including responses to antibiotic stress (Moreno-Gámez et al., 2017). Competence activation can also alter pneumococcal survival during antibiotic exposure, with effects that depend on the antibiotic (Prudhomme et al., 2024).

Studies of *luxS* mutants have demonstrated effects on this process. Trappetti et al. (2011) reported reduced transformation efficiency in a LuxS-deficient pneumococcal strain. Romão et al. (2006) found that *luxS* inactivation altered the timing and regulation of competence, and that culture supernatants from *luxS*-positive, CSP-export-defective pneumococci could complement the competence phenotype. These experiments support the involvement of an extracellular factor, but the presence of other compounds in conditioned supernatants makes it difficult to attribute the effects specifically to AI-2. DPD supplementation therefore provides a more specific approach to examining the contribution of the AI-2 precursor to transformation.

Here, we investigated the contribution of LuxS to pneumococcal responses to ampicillin and erythromycin and examined whether DPD supplementation could alleviate the antibiotic-associated phenotype of a Δ*luxS* mutant. We then assessed the effect of *luxS* inactivation on genetic transformation and tested whether DPD supplementation could recover transformation efficiency. Together, these experiments address the contribution of DPD/AI-2 to antibiotic stress responses and genetic competence in *S. pneumoniae*.

## MATERIALS AND METHODS

### Bacterial strains and media

All strains of *S. pneumoniae* used are listed in Table 1. Streptococci were stored at −80 °C in Todd Hewitt Broth (THB, Becton Dickinson and Company, Le Pont de Claix, France) or Tryptic Soy Broth (TSB, Soybean-Casein Digest medium, Bacto™) supplemented with 30% glycerol. Precultures used in the experiments described below were made from fresh liquid cultures incubated at 37 °C in a 5% CO2-supplemented atmosphere and grown until the cultures reached an absorbance at 600 nm (optical density at 600 nm [OD600]; Biophotometer; Eppendorf) of 0.3–0.5, before storage at −80 °C in 15% glycerol. For transformation assays, semi-defined media C+Y_YB_ (Stevens et al., 2011; Salvadori et al., 2016) was used. For plating, blood agar base No. 2 (Oxoid, Hampshire, England) supplemented with 5% defibrinated sheep blood (TCS Biosciences Ltd., Buckingham, United Kingdom) was used. Final concentrations of selective levels of antibiotics were 500 µg mL−1 kanamycin and 200 µg mL^−1^ spectinomycin.

**Table 1.**
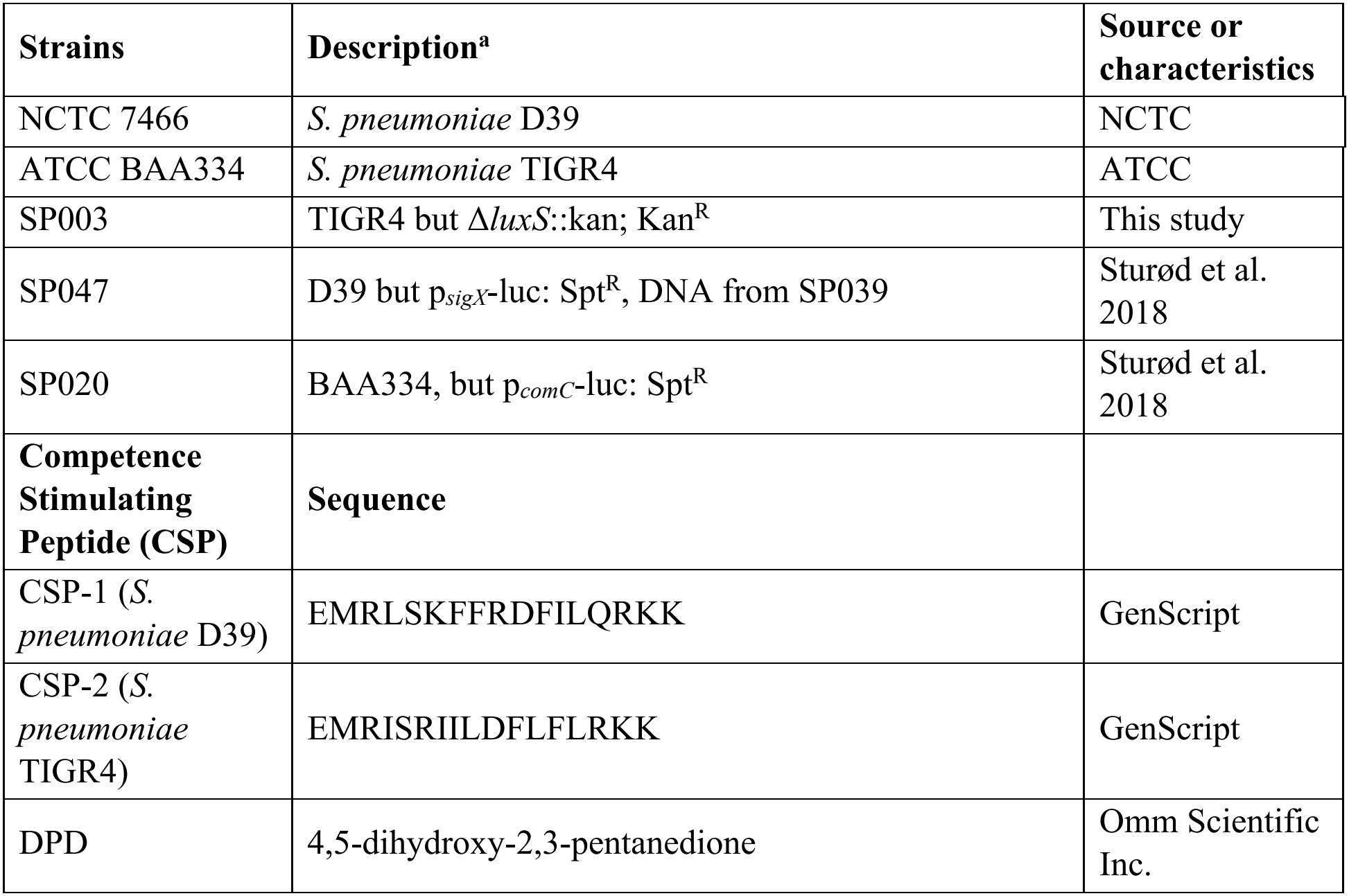
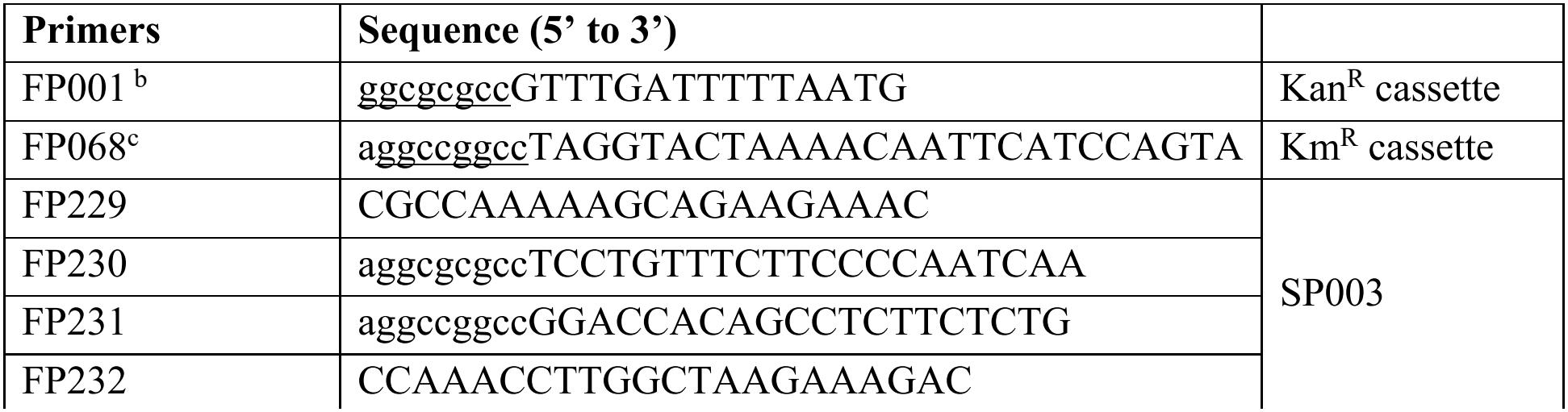
Strains, mutants, synthetic peptides, molecules and primers used in this study.

### Synthetic peptide, DPD and antibiotics

The sequence of the synthetic CSPs used in this study is listed in Table 1. Stock solutions were prepared by resuspending CSP-1 and CSP-2 in distilled water to a concentration of 0.5 mM and 10 mM, respectively, and storing it at −20 °C. For each experiment, antibiotics were diluted into pre-defined concentrations in C+Y_YB_. Antibiotic stocks were prepared in distilled water (erythromycin was prepared in ethanol) and stored at −20 °C at the following concentrations: ampicillin (50 mg mL^−1^) and erythromycin (20 mg mL^−1^). DPD was obtained from Omm Scientific Inc. (Texas, USA). A 7.5 mM DPD stock solution was prepared in sterile ultrapure water and stored at −20 °C.

### Preparation of genomic donor DNA

Genomic donor DNA was extracted from strains according to the CTAB method previously described(1).

Construction of the Δ***luxS*** mutant. The TIGR4 Δ*luxS* mutant SP003 was constructed by PCR ligation mutagenesis (Lau et al., 2002; Petersen et al., 2005). Regions upstream (625 bp) and downstream (631 bp) of *luxS* were amplified using primer pairs FP229–FP230 and FP231– FP232, respectively. The kanamycin-resistance cassette was amplified using FP001–FP068 (Sung et al., 2001). AscI restriction sites were introduced through FP230 and FP001, and FseI sites through FP231 and FP068. Following digestion with the corresponding restriction enzymes, the upstream and downstream fragments were ligated separately to the kanamycin-resistance cassette using T4 DNA ligase (Fermentas). The two ligation products were mixed and amplified using FP229–FP232 to generate the complete replacement construct, which was purified using the QIAquick PCR purification kit (Qiagen).

For transformation, precultures at OD_600_ 0.5 were diluted 1:10 in TSB and incubated at 37 °C with 5% CO_2_ until OD_600_ reached 0.1. CSP-2 was added at a final concentration of 250 nM together with the purified construct. Cultures were incubated aerobically at 37 °C for 4 h and plated on blood agar containing kanamycin. Transformants were recovered after 24 h at 37 °C with 5% CO_2_. Replacement of *luxS* with the kanamycin-resistance cassette was checked by PCR using genomic DNA from the mutant and primer pair FP229–FP232 to assess the size of the amplified region.

Antibiotic exposure and DPD supplementation. TIGR4 and its isogenic Δ*luxS* mutant SP003 were inoculated into TSB at approximately 7 × 10^6^ CFU mL^−1^. Cultures containing erythromycin (0.02–2 mg L^−1^) or ampicillin (0.02–0.1 mg L^−1^), with or without DPD supplementation, were dispensed into 96-well microtiter plates. DPD was tested at concentrations ranging from 30 to 200 nM. Cultures without antibiotics were included as controls. Each well contained 200 µl of bacterial culture and plates were incubated at 37 °C AIR. Each well contained 200 µl of bacterial culture and plates were incubated at 37 °C AIR. After 6 h, cultures were serially diluted and plated on blood agar. Plates were incubated at 37 °C with 5% CO_2_ for 24 h, after which viable counts were determined and expressed as CFU mL^−1^.

Transformation-efficiency assays. Transformation assays were performed using the procedure described by Salvadori et al. (2016), with the conditions specified below. Precultures of TIGR4 and SP003 at OD_600_ 0.5 were diluted 1:100 in C+Y_YB_ and incubated at 37 °C with 5% CO_2_ for 15 min. Competence was induced by adding CSP-2 at a final concentration of 250 nM together with a spectinomycin-resistance genomic donor DNA from SP020. DPD was added right after CSP addition atfinal concentrations of 500 nM, 2 µM and 4 µM, alongside controls without added DPD.

Cultures were incubated at 37 °C for 4 h. Appropriate dilutions were plated on blood agar with and without spectinomycin to determine transformant counts and total viable counts, respectively. Plates were incubated at 37 °C with 5% CO_2_ for 24 h. Transformation efficiency was calculated as the number of spectinomycin-resistant CFU divided by the total viable CFU, multiplied by 100, and expressed as a percentage.

### Luciferase reporter assays

Promoter activity was assessed using SP047, carrying the PsigX-luc reporter in the D39 background, and SP020, carrying the PcomCDE-luc reporter in the TIGR4 background. Fresh colonies from blood agar were inoculated into C+YYB containing DPD at concentrations of 250 nM–8 µM. Assays were performed with or without the corresponding CSP: CSP-1 for SP047 and CSP-2 for SP020, at a final concentration of 50 nM.

Culture aliquots of 200 µL were dispensed into transparent-bottom 96-well microtiter plates (Nunc, Thermo Scientific). Uninoculated medium was included as a blank. D-Luciferin (Synchem, Felsberg-Altenberg, Germany) was added to each well as 10 µL of a 1 mM solution. Plates were incubated at 37 °C in air. Relative luminescence units (RLU) and OD_600_ were recorded at 15-min intervals during growth using a Synergy HT microplate reader (BioTek, Winooski, Vermont, USA). Luminescence was normalized to the corresponding OD_600_ and expressed as RLU/OD_600_.

### Statistical analysis

Data from three independent biological experiments were analyzed, with technical replicates averaged within each experiment. CFU counts and transformation efficiencies were log_10_-transformed when appropriate. Differences between experimental conditions were assessed using two-sided paired Student’s t-tests. The Holm method was applied to correct for multiple comparisons, and adjusted p values below 0.05 were considered statistically significant.

## Results

### Impact of DPD concentrations on the growth of *S. pneumoniae* and its Δ*luxS* mutant

DPD concentrations of 4 and 10 µM had little apparent effect on planktonic growth of TIGR4 or Δ*luxS* SP003, whereas 50 and 100 µM inhibited growth in representative cultures (Fig. 1). Subsequent assays therefore used DPD concentrations of 8 µM or lower.

**Figure 1.**
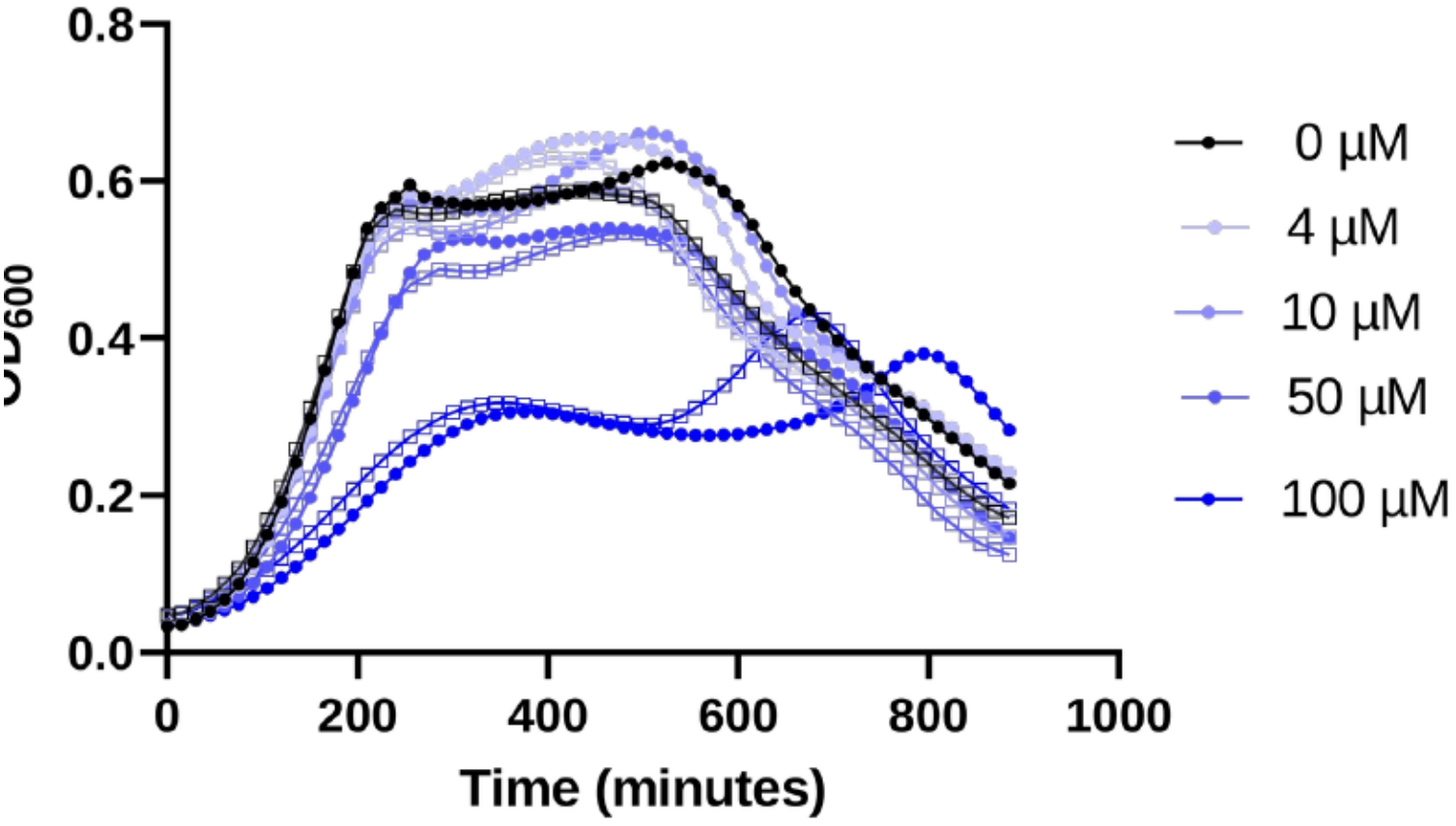
Effect on growth of different DPD concentrations in *S. pneumoniae* TIGR4 (filled circles) and its isogenic *luxS* knockout (open squares) strain. Fresh colonies of the two strains were grown in C+Y_YB_ and aliquots were placed on a 96-well plate with different concentrations of DPD. Culture growth was measured every 15 minutes during incubation at 37 °C in 5% CO_2_ atmosphere. Growth was quantified using the area under the OD600 growth curve (AUC). Statistical significance was assessed by two-way repeated-measures ANOVA, with strain and DPD concentration as factors. These values are presented as averages from three replicates and are representative of three independent experiments.

### Inactivation of *luxS* in *S. pneumoniae* increases antibiotic susceptibility

To assess the effect of *luxS* inactivation on antibiotic susceptibility, TIGR4 and its isogenic Δ*luxS* mutant SP003 were exposed to erythromycin or ampicillin for 6 h before CFU enumeration (Fig. 2). Because untreated CFU differed between strains, antibiotic responses were additionally analyzed as log10 reductions relative to the no-antibiotic control for each strain within each biological experiment. Across the tested concentrations, the mutant showed greater relative reductions in CFU than WT for both antibiotics (Holm-adjusted p < 0.05), indicating that the difference in antibiotic response was not explained solely by baseline CFU differences.

**Figure 2.**
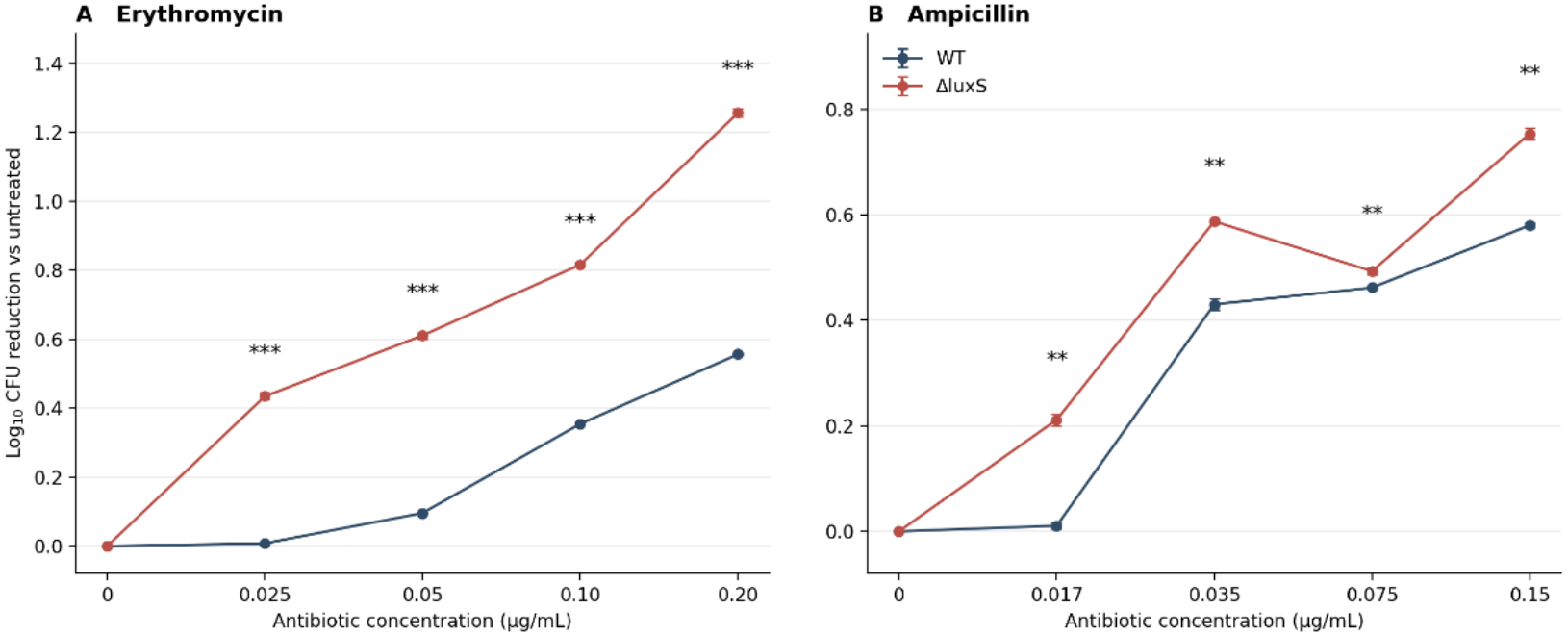
Effect of erythromycin and ampicillin on the viability of *S. pneumoniae* TIGR4 and its isogenic Δ*luxS* mutant SP003. Strains were exposed to the indicated concentrations of (A) erythromycin or (B) ampicillin for 6 h. Viable counts were determined by plating, and log_10_ CFU reductions were calculated relative to the untreated control of each strain within each experiment. Data represent mean ± SD from three independent biological experiments. Statistical differences between WT and Δ*luxS* were assessed using two-sided paired *t*-tests with Holm correction (**adjusted *p* < 0.01; ***adjusted *p* < 0.001).

### DPD supplementation influences antibiotic-associated viability of the *luxS* mutant

DPD supplementation at 50 nM increased viable counts of Δ*luxS* during exposure to both antibiotics (Fig. 3). Across three independent experiments, CFU increased 3.54-fold with erythromycin (Holm-adjusted p = 0.0264) and 2.86-fold with ampicillin (adjusted p = 0.0019), relative to the corresponding unsupplemented mutant. At 200 nM, responses varied among experiments and did not reach statistical significance after correction (adjusted p = 0.5201 for both antibiotics). Shared control measurements in the original data were used as the corresponding reference without treating repeated entries as additional independent observations.

**Figure 3.**
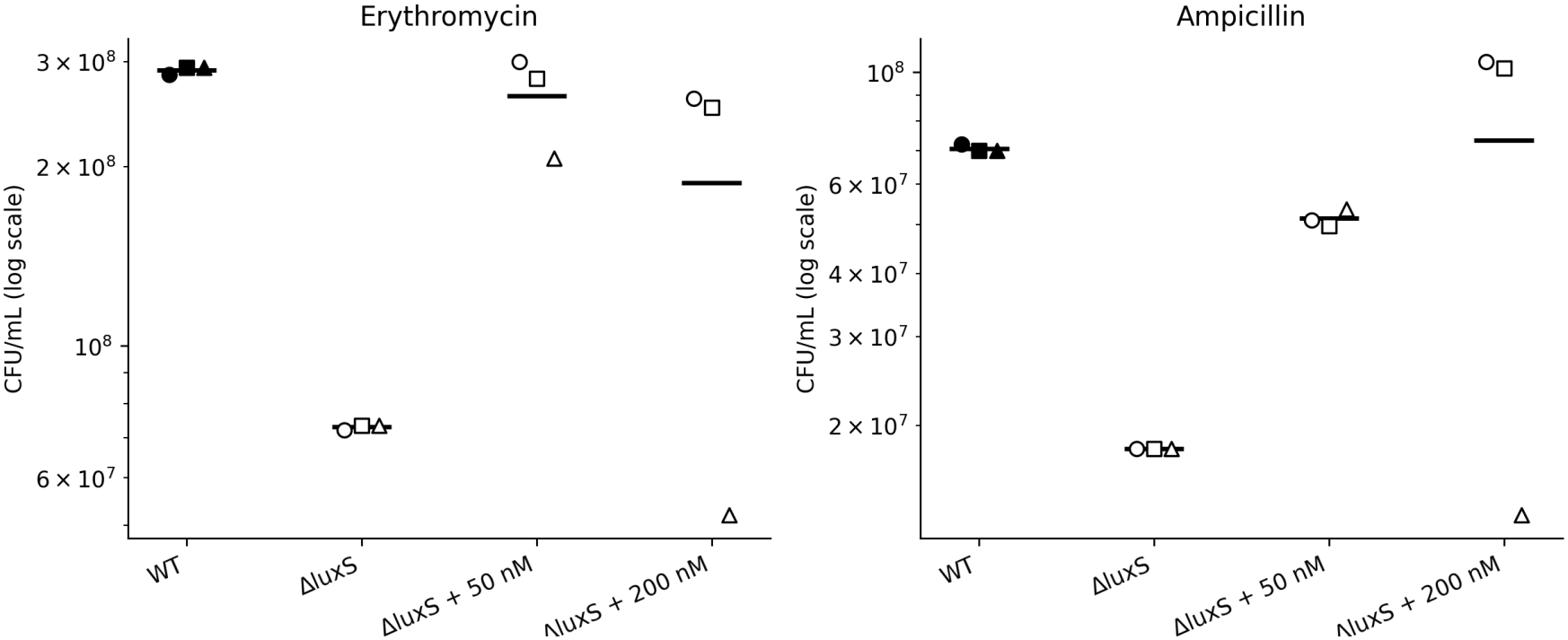
Effect of DPD supplementation on viable counts of *S. pneumoniae* TIGR4 and Δ*luxS* following 6 h exposure to erythromycin or ampicillin. The principal comparisons were Δ*luxS* with 50 or 200 nM DPD versus unsupplemented Δ*luxS*. Data are from three independent biological experiments; technical measurements were averaged within experiments. Log_10_ ratios were evaluated with two-sided one-sample t-tests and Holm correction across four planned comparisons. Supplementation with 50 nM DPD increased viable counts significantly under both antibiotic exposures (adjusted p < 0.05), whereas the effect of 200 nM DPD was not significant.

### Effect of DPD on competence-associated promoter activity

We monitored CSP-responsive *sigX* and *comCDE* promoter activity using luciferase reporters during DPD supplementation (Fig. 4). In the representative time courses, DPD was associated with reduced CSP-associated *sigX* reporter output, whereas the corresponding effect on *comCDE* was less evident. Reporter measurements are described qualitatively, without treating repeated time points as independent replicates.

**Figure 4.**
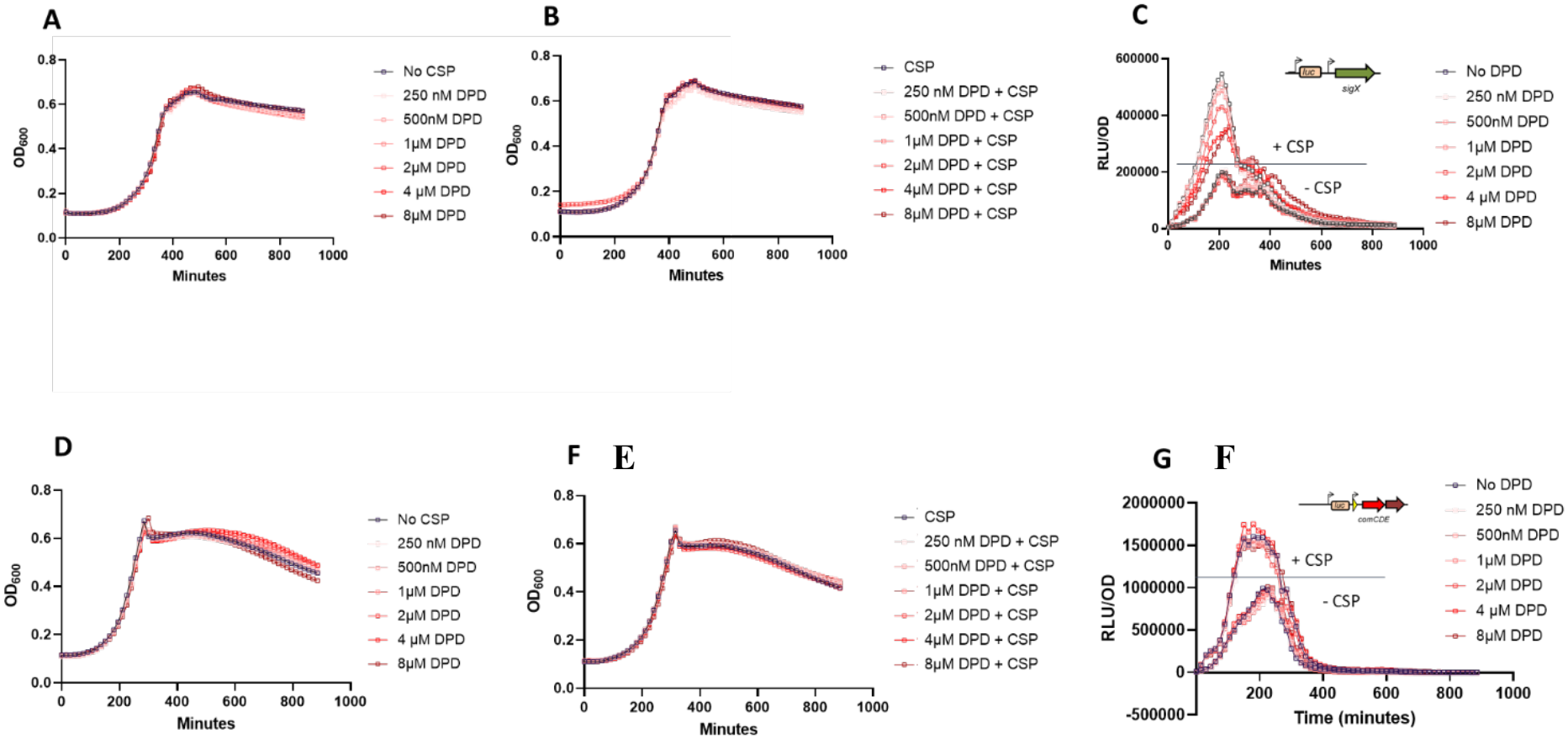
Effect of DPD supplementation on pneumococcal growth and competence-associated promoter activity. SP047, carrying the PsigX-luc reporter in the D39 background (A–C), and SP020, carrying the PcomCDE-luc reporter in the TIGR4 background (D–F), were grown in C+YYB with the indicated DPD concentrations. Growth was monitored by OD_600_ in the absence (A, D) or presence (B, E) of CSP. Promoter activity is presented as luminescence normalized to OD_600_ (C, F). CSP-1 was used for SP047 and CSP-2 for SP020 at 50 nM. Curves show replicate cultures from a representative experiment; three independent experiments were performed.

### DPD supplementation partially restores transformation efficiency in the *luxS* mutant

Across three independent experiments, transformation efficiency in untreated Δ*luxS* was 0.23-fold that in untreated WT (geometric mean ratio; Holm-adjusted p = 0.013). Supplementation of Δ*luxS* with 0.5, 2, or 4 µM DPD increased its transformation efficiency by 2.14-, 2.52-, and 2.75-fold, respectively, relative to untreated Δ*luxS* (Holm-adjusted p = 0.013, 0.0092, and 0.0059; Fig. 5). Nevertheless, each supplemented mutant group remained significantly below the untreated WT (geometric mean ratios 0.49, 0.58, and 0.63; adjusted p = 0.013, 0.018, and 0.018, respectively). These paired log-ratio analyses indicate a significant but partial rescue of transformation efficiency by DPD.

**Figure 5.**
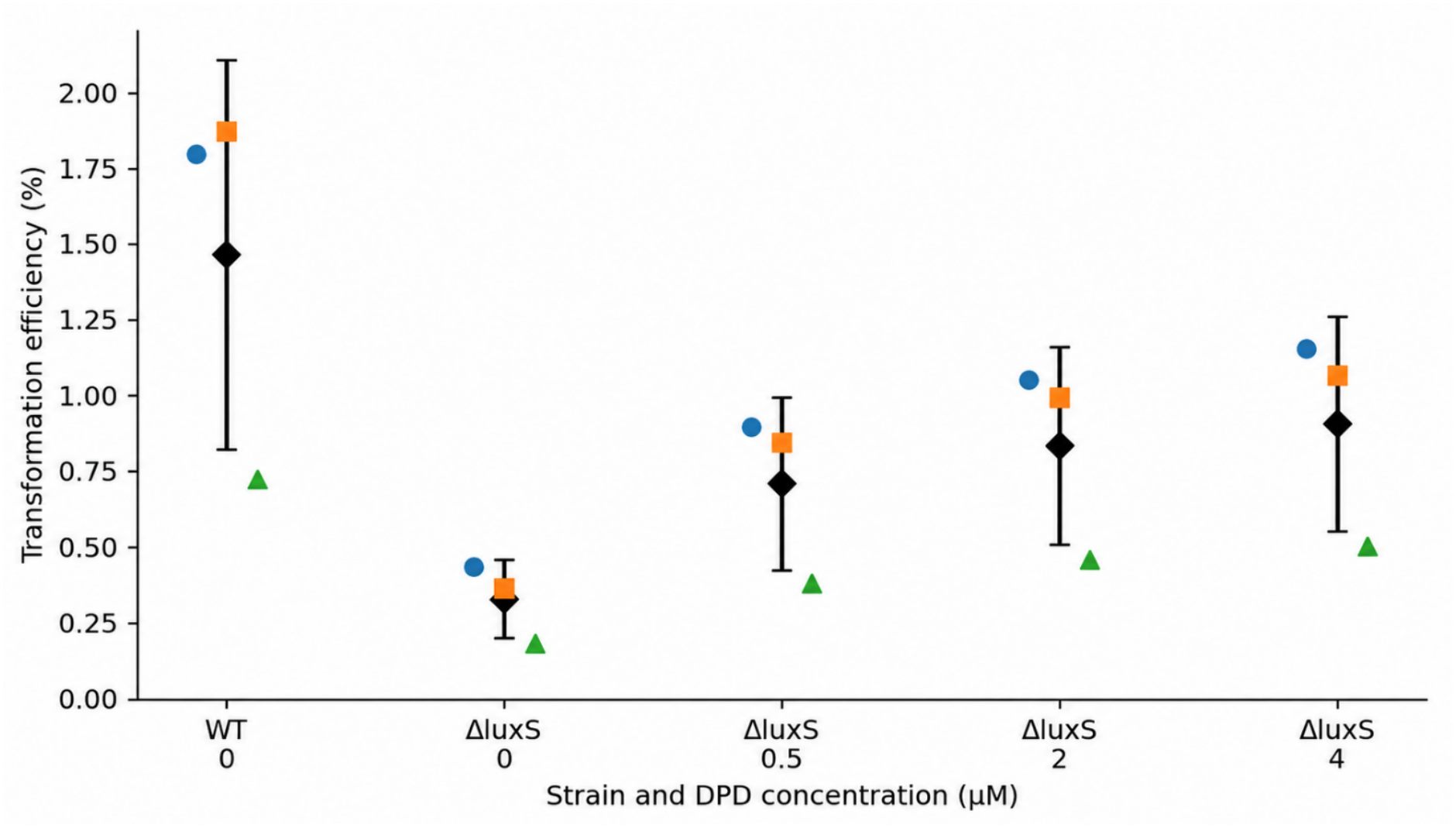
DPD supplementation partially restores transformation efficiency of *S. pneumoniae* Δ*luxS*. Untreated WT and Δ*luxS* without DPD or with 0.5, 2, or 4 µM DPD were analyzed in three independent biological experiments, each with three technical measurements. Experiment-level means were compared using paired log_10_ ratios and two-sided one-sample t-tests with Holm correction across seven planned contrasts. The displayed groups focus on the deletion phenotype and its response to DPD.

## Discussion

Our findings support a role for LuxS in pneumococcal antibiotic responses and genetic competence. Relative to their strain-matched untreated controls, Δ*luxS* cultures exhibited larger antibiotic-associated reductions in CFU than the wild type. Supplementation with 50 nM DPD significantly increased mutant CFU during exposure to both antibiotics, whereas the response to 200 nM was variable. DPD also partially restored transformation efficiency in the mutant.

Previous work in *S. pneumoniae* showed that exposure to subinhibitory concentrations of penicillin induces *luxS* expression (Rogers et al., 2007), suggesting a possible involvement of LuxS in the pneumococcal response to antibiotic stress. Our study provides functional evidence supporting this association by demonstrating increased antibiotic susceptibility following *luxS* inactivation. Importantly, the greater reduction in viable counts in the mutant persisted after accounting for differences in baseline growth, and supplementation with DPD partially alleviated this phenotype. These findings suggest that LuxS-associated processes contribute to pneumococcal survival during antibiotic exposure, although the underlying mechanisms remain to be elucidated.

The effects of DPD also appear to be concentration-dependent. While low concentrations partially alleviated the antibiotic susceptibility phenotype of the Δ*luxS* mutant, higher concentrations inhibited bacterial growth, consistent with previous observations in pneumococcal clinical isolates (Agnew et al., 2023). These findings highlight the importance of DPD concentration and experimental conditions when interpreting LuxS/AI-2-associated phenotypes.

The involvement of LuxS in pneumococcal genetic competence has also been documented (Romão et al., 2006; Trappetti et al., 2011). However, whether exogenous DPD can compensate for the reduced transformation efficiency associated with *luxS* inactivation has remained unclear. Here, we demonstrate that supplementation with defined concentrations of DPD significantly increases transformation efficiency in the Δ*luxS* mutant, although it does not fully restore wild-type levels. This finding extends previous observations by demonstrating that DPD supplementation can partially compensate for the loss of LuxS activity in pneumococcal transformation.

Because LuxS participates in both DPD production and the activated methyl cycle, the phenotypes associated with its inactivation cannot be attributed exclusively to AI-2 signaling. Nevertheless, the partial restoration of survival during antibiotic exposure and genetic transformation following DPD supplementation supports a contribution of DPD-associated processes to both phenotypes. Further studies will be needed to determine the underlying mechanisms and distinguish the metabolic and signaling contributions of LuxS.

Together, our findings establish a role for LuxS in pneumococcal survival during antibiotic exposure and provide new evidence that exogenous DPD can partially compensate for the effects of *luxS* inactivation on both antibiotic susceptibility and genetic transformation. These results extend our understanding of LuxS-associated phenotypes in *S. pneumoniae* and highlight the importance of distinguishing the metabolic and signaling contributions of this pathway. Further investigation of the mechanisms underlying these effects may reveal how LuxS-associated processes influence antibiotic responses and the acquisition of genetic material in pneumococci.

## Acknowledgements

This work is part of the Project Internet of Bio-Nano Things for Prediction and Prevention of Infectious Diseases funded by the Research Council of Norway Project nr. 310601.

## Notes

### Competing Interest Statement

The authors have declared no competing interest.

